# Microexon alternative splicing and feeding behavior in *C. elegans*

**DOI:** 10.64898/2026.05.26.727998

**Authors:** Gonzalo Monti, Diego Rayes, Alberto R. Kornblihtt, Micaela A. Godoy Herz

## Abstract

Microexons are short alternative exons up to 51 nucleotides long that are highly enriched in neuronal genes. Their dysregulation has been linked to human neurodevelopmental disorders, including autism spectrum disorders. In the nematode C*aenorhabditis elegans*, global regulation of microexons is also critical for proper development. Here we show that microexon alternative splicing (AS) changes between *C. elegans* larval and adult stages and that microexon inclusion is differentially regulated among distinct neuronal types. Consistently with previous evidence that *C. elegans* splicing is regulated in response to environmental stimuli, we found here that specific microexons are modulated upon food availability. Both the inclusion levels of these microexons and the feeding behavior seem to depend on the DNA topology, which may affect transcription dynamics, as revealed by the effects of the topoisomerase I (TOP1) inhibitor, camptothecin (CPT). CPT treatment alters responses related to food availability such as speed reduction and exploration. Furthermore, animals carrying a mutation in the global regulator of microexon splicing *prp-40* exhibit altered food preference, independently demonstrating that disruption of microexon AS has important consequences on animal behavior.

## Introduction

Alternative splicing (AS) is one of the key steps in RNA processing that contributes to diversity in transcriptomes: various mature messenger RNAs (mRNAs) can be obtained from a single gene (Ast 2004; House and Lynch 2008; Rogalska et al. 2023).

AS plays a crucial role at both the cellular and organismal levels. It contributes to development in multiple ways and modulates responses to environmental stimuli (Ip et al. 2011; Godoy Herz et al. 2019; Marasco et al. 2022). The most common type of AS event in both humans and *C. elegans* is cassette exon inclusion, a mode in which transcript isoforms differ in whether a particular exon is included or skipped. A subset of these events involves microexons, extremely short alternative exons, typically shorter than 27 nucleotides. In this work we will refer to microexons as exons shorter than 51 nucleotides (Li et al. 2015). Microexons are abundant in genes expressed in neurons and represent the most evolutionarily conserved AS events in vertebrates (Torres-Méndez et al. 2019). The inclusion of microexons is thought to modulate the function of interaction domains of proteins involved in neurogenesis where the interaction surface seems to be more important than the microexon sequence itself (Irimia et al. 2014). In addition, the inclusion of microexons is under strong selective pressure throughout nervous system development (Irimia et al. 2014). The relevance of this process is evident, as it has been suggested that its dysregulation may underlie human neurodevelopmental disorders such as autism spectrum disorders (ASD) (Irimia et al. 2014). Apart from ASD, patients with other neuronal disorders such as schizophrenia and epilepsy have been shown to exhibit dysregulation of various microexon events (Gandal et al. 2018; Gonatopoulos-Pournatzis et al. 2020). Nevertheless, the functions and regulatory mechanisms of most human-relevant microexons remain unknown.

Elucidating these fundamental processes requires *in vivo* models where splicing events can be linked to specific behavioral and developmental outputs. Recently it has been shown that global regulation of microexons also profoundly affects the development of the nematode *C. elegans* (Choudhary et al. 2021). The authors reported that the spliceosomal protein PRP-40 (the human ortholog of PRPF40A) specifically regulates microexon inclusion at a genome-wide level in *C. elegans*. Worms carrying mutations in the *prp-40* gene exhibit severe developmental defects and die before reaching adulthood. Notably, many genes containing PRP-40–regulated microexons have human orthologs associated with neurodevelopmental disorders such as ASD.

A genome-wide RNA-seq analysis of *prp-40* mutants showed that, in the absence of PRP-40, microexons are preferentially excluded compared with wild-type animals, indicating that *prp-40* is required for their proper inclusion (Choudhary et al. 2021). However, the function of most microexons remains unknown. Only recently have the roles of specific *C. elegans* microexons been investigated, and a small number of these are starting to be functionally characterized (Choudhary et al. 2025).

In this study we aimed to investigate the role of microexons in *C. elegans*, focusing on their regulation during development and in response to an environmental stimulus, specifically food availability.

We found that microexon alternative splicing is regulated between larvae and adults, and that the inclusion of a given microexon varies among different neuronal types. Furthermore, microexon alternative splicing is modulated upon food availability in a subset of genes. Finally, we report a role for DNA topology in this process, revealed by alterations in both microexon alternative splicing and feeding behavior caused by inhibition of topoisomerase I.

## Results

### Microexon AS changes between *C. elegans* larvae and adults

Based on RNA-seq data from the *prp-40* mutant (Choudhary et al. 2021), we selected a panel of AS events regulated by PRP-40 to assess changes in development. For this selection, we focused on genes that at the same time are known to be relevant for neuronal functioning and exhibit the strongest changes in microexon inclusion when comparing the *prp-40* mutant strain to wild-type animals. *Prp-40* mutant animals arrest their development at the larval stage 3 (L3), indicating that proper regulation of microexon splicing is essential for development. Accordingly, we first asked whether microexon inclusion changes in the selected genes between total RNA samples from first larval stage (L1) and adult animals. To address this, we examined microexon AS in endogenous genes in wild-type worms using RT-PCR followed by polyacrylamide gel electrophoresis. This approach allows us to distinguish the isoform including the microexon from the one that excludes it, which, as noted, differ by only a few nucleotides. Changes in AS were quantified using the Splicing Index, defined as the ratio of the longest isoform to the total isoforms.

We observed that microexon inclusion levels are deeply affected when comparing larval and adult stages. These changes show distinct trends across the different events studied: in some cases, microexon inclusion increases from larvae to adults, as seen for *kvs-5,* encoding a potassium channel, whereas in others it decreases during development, as illustrated by *tyra-2, lgc-40,* or *rgr-1,* encoding a tyramine receptor, a ligand-gated ion channel and a component of the mediator complex, respectively (Figure 1). We included the 84-nt cassette exon of *unc-16* because, although it is longer than 51-nt, it is also regulated by *prp-40* (Choudhary et al. 2021). We also included the glutamate receptor *glr-4* gene that showed no changes and therefore functions as a not-responding control. These results suggest that microexon AS is developmentally regulated, and that this regulation is not uniform across all microexons.

**Figure 1:**
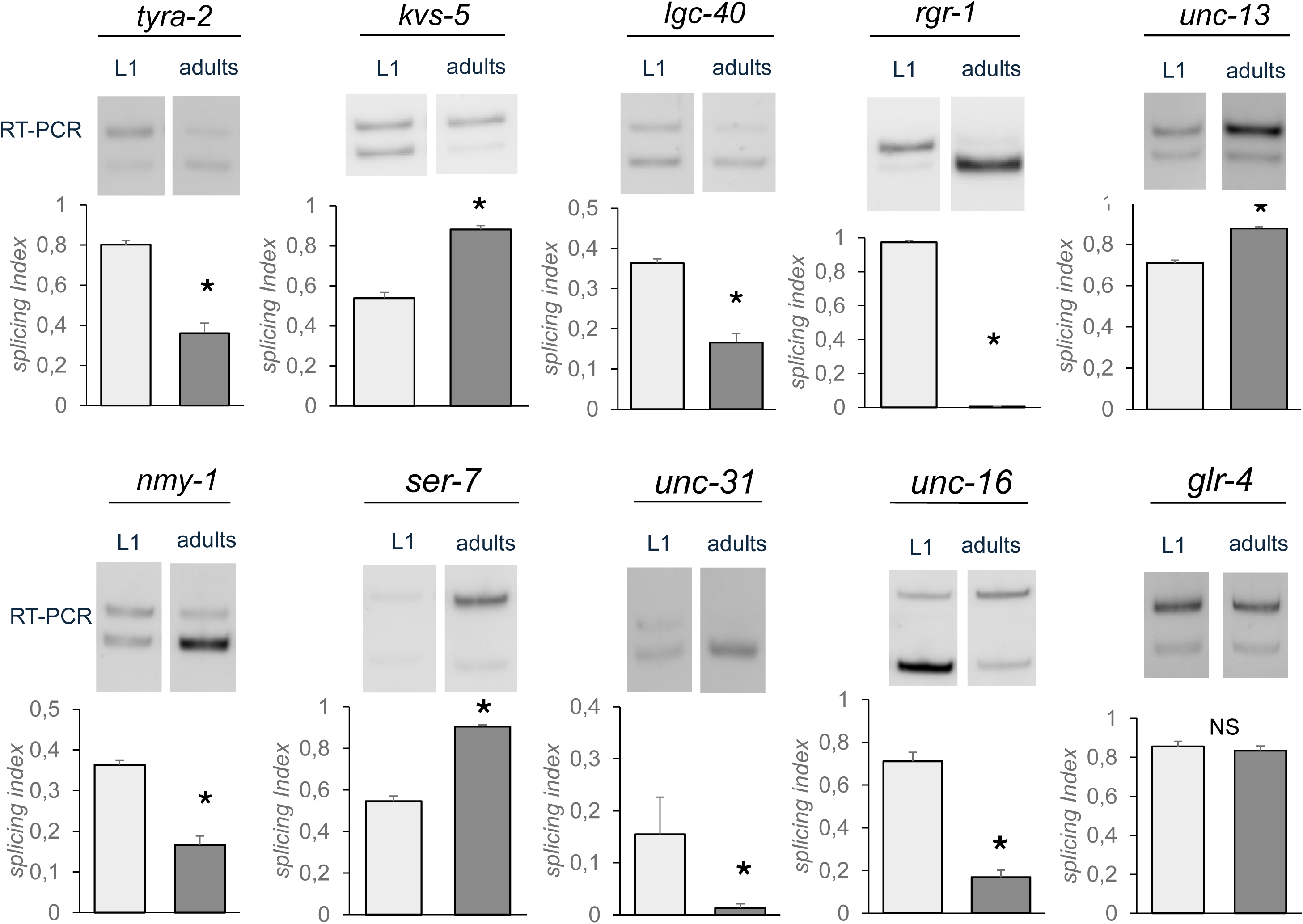
Microexon AS changes between *C. elegans* larvae and adults. Top: representative RT-PCR gel images for the alternative splicing patterns of microexons, from left to right: *tyra-2, kvs-5, lgc-40, rgr-1, nmy-1, ser-7, unc-31* and *glr-4*. Bottom: quantification of splicing index (SI) is calculated as the amount of the longest isoform relative to the amount of all isoforms. White bars, L1; gray bars, adults. Data represent means ± SD (n ≥ 3). **\*** p < 0.05 (Student’s t-test).

We also examined what occurs at the L3 stage, an intermediate developmental stage between L1 and adulthood, given that *prp-40* mutant worms survive only until this stage. We performed the experiment described in Figure 1 and, after RNA extraction, analyzed alternative splicing changes by RT-PCR in animals at the L1, L3, and adult stages. In the cases of *rgr-1* and *unc-16*, we observed intermediate levels of microexon inclusion at L3 (Supp. Fig. 1).

### Microexon inclusion is differentially regulated among distinct neuronal types

In order to investigate if regulation of microexon alternative splicing is important at the spatial level, we used available nervous system–specific RNA-seq data (Taylor et al. 2021; Weinreb et al. 2025) to assess microexon inclusion across different neuronal types. We observed that microexon inclusion profiles vary considerably depending on the neuronal type, and that each microexon exhibits a distinct pattern (Figure 2 A-C), suggesting that regulation is not uniform across all microexons studied.

**Figure 2:**
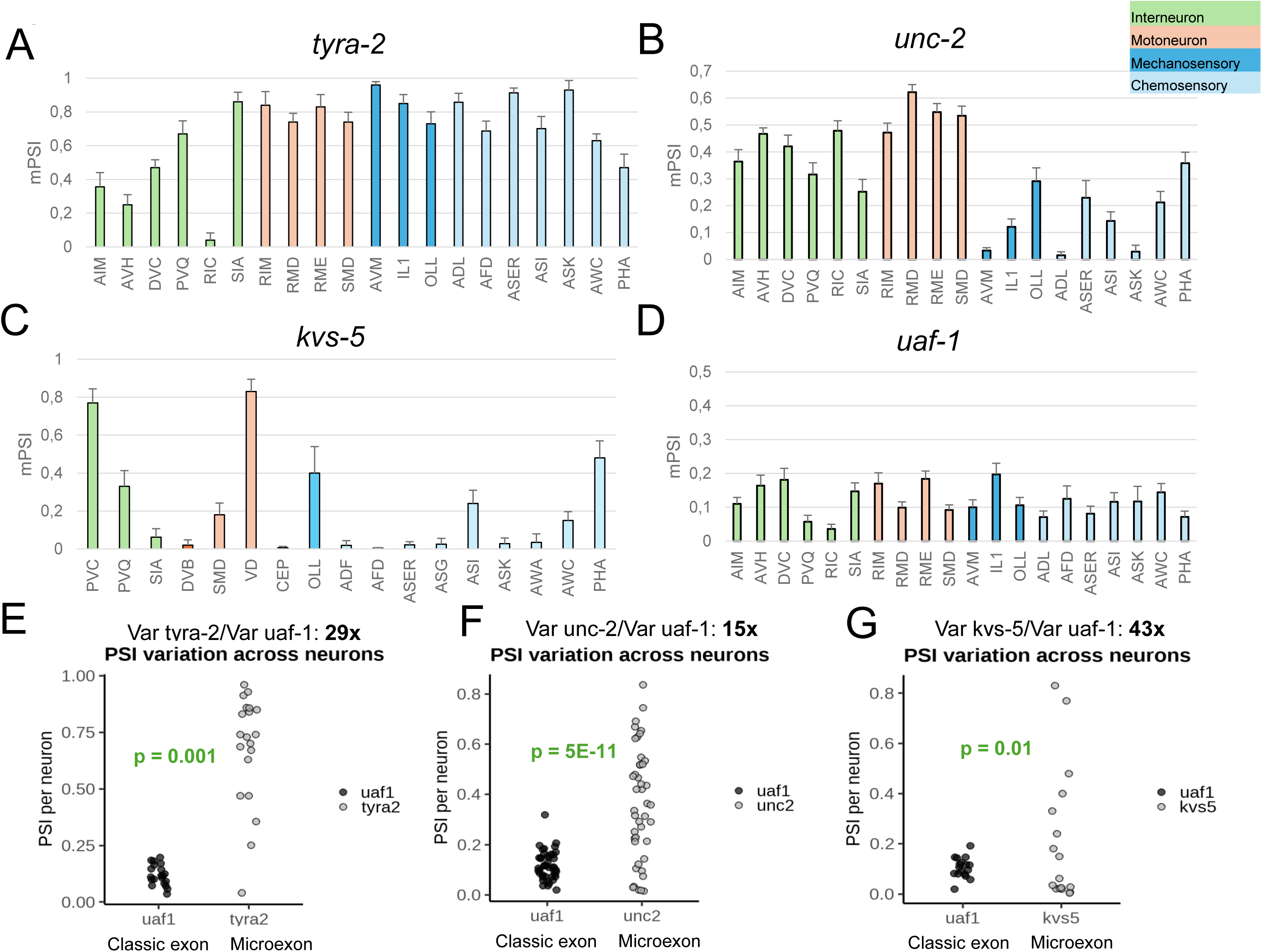
Microexon inclusion is differentially regulated among distinct neuronal types. Analysis of neuron-specific microexon AS (Taylor et al, 2021). Mean Percent Selected Index (mPSI) for each neuronal type for microexons *tyra-2* (A), *unc-2* (B) and *kvs-5* (C) and for classic exon *uaf-1* (D). Green bars, interneurons; orange bars, motoneurons; dark blue, mechanosensory neurons; light blue, chemosensory neurons. Mean and SD are shown. Comparison of variances of mPSI between each microexon and the control exon *uaf-1* for *tyra-2* (E), *unc-2* (F) and *kvs-5* (G). Fold change between variances and their significant p-values (Levene’s test) are shown.

As a control with a non-microexon AS event, we examined the inclusion levels of the 216 nt-long cassette exon of the *uaf-1* gene, which resulted less variable across neuron types than the microexons (Figure 2D). *Uaf-1* encodes a splicing regulator. A statistical comparison of variances between the microexons *tyra-2*, *unc-2* and *kvs-5* and the *uaf-1* non-microexon control revealed a significant difference (Figure 2 E-G). This coincides with the results of Weinreb at al. (2025) that show that microexons display more variable AS inclusion profiles than non-microexons, demonstrating that AS is spatially regulated within the nervous system.

### Microexon AS changes upon food availability

In addition, we asked whether, beyond the developmental changes observed, *C. elegans* microexon AS is modulated by environmental stimuli. We chose food availability as the stimulus. Previous work had shown that *C. elegans* AS is altered upon food availability during larval stages (Baugh et al. 2009), without a particular analysis on the specific impact on microexon AS.

To address this, we designed the following experiment: adult animals were transferred to plates without food (starvation conditions) for two hours. They were then moved to either plates containing food (*E. coli* OP50 strain) or plates lacking it. Since worms avoid contacts with copper, we placed a copper ring on the medium surface to limit the field of exploration and prevent the worms from escaping during the starvation period (Davies et al. 2003). After two hours on the new plates, animals were harvested, total RNA was extracted, and microexon AS isoforms were assessed by RT-PCR (Fig. 3A).

**Figure 3:**
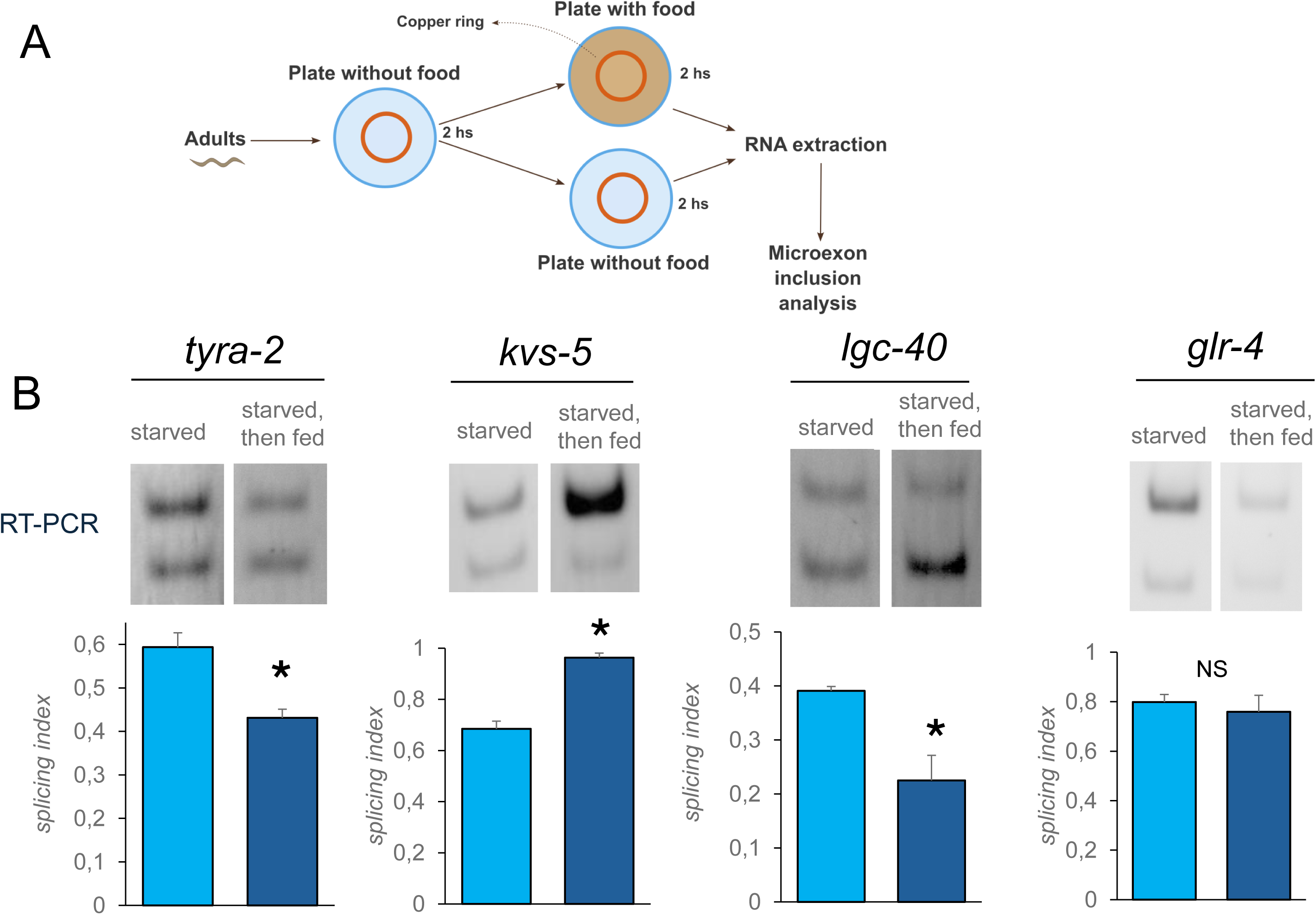
Microexon AS changes upon food availability. (A) Food availability protocol. (B) Top: Representative RT-PCR gel images for the alternative splicing patterns of *tyra-2, kvs-5, lgc-40* and *glr-4*. Bottom: quantification of SI. Light blue, starved worms; dark blue, starved then fed worms. Data represent means ± SD (n ≥ 3). **\*** p < 0.05 (Student’s t-test).

We observed changes in microexon inclusion in *kvs-5*, *tyra-2*, and *lgc-40* when comparing starved and fed worms. In *tyra-2* and *lgc-40*, microexon inclusion decreased, whereas in *kvs-5* it increased when worms were re-seeded on food-containing plates. As a control, the *glr-4* gene showed no changes (Fig. 3B), consistently with its insensitivity to developmental changes (Fig. 1 and Suppl. Fig. 1). *Tyra-2* encodes a tyramine receptor, a neurotransmitter associated with the response to food encounter in *C. elegans*. *Kvs-5* encodes a potassium channel and *lgc-40* encodes an ion channel that responds to serotonin. Interestingly, serotonin is a neurotransmitter that modulates the response to feeding in *C. elegans*, and its effect is the opposite to tyramine.

These results indicate that AS of specific microexons is modulated in response to an environmental stimulus such as food availability. Notably, both *tyra-2* and *lgc-40* are important receptors in food-related behaviors (Fu et al. 2018).

### Mutation of the microexon regulator *prp-40* alters food preference behavior

Given that microexon AS changes in response to food availability, we asked whether animals carrying a mutation in *prp-40*—the regulator of microexon splicing with globally altered AS—also exhibit changes in behavior upon encountering food. In wild-type worms, it is well established that starved animals reduce their locomotion and spend more time feeding when they encounter food, compared to well-fed (satiated) animals (Sawin et al. 2000; Flavell et al. 2013). We sought to determine how this behavior is affected in the *prp-40* mutant strain. We first noted that *prp-40* animals display locomotor defects, as reported previously (Choudhary et al. 2021). Therefore, prior to conducting behavioral assays, we performed a locomotion assay as a prerequisite control. Specifically, we carried out a thrashing assay, which quantifies the number of body bends (thrashes) per unit time and found that the number of thrashes per minute was markedly reduced in *prp-40* mutants compared to wild-type animals (Figure 4A). This confirms that the mutants exhibit locomotor defects, which makes it unsuitable to perform a standard food encounter behavioral assay, as their basal movement is already severely impaired. Upon this impediment, we opted for a more refined behavioral paradigm, aimed at assessing decision-making rather than locomotion *per se*. Specifically, we examined whether *prp-40* mutant animals display a differential behavior when presented with a choice between two distinct food sources. To this end, we performed the experiment shown in Fig. 4B: wild-type and *prp-40* mutant animals, previously grown on a lawn of OP50 *E. coli*, were transferred to a new plate where they could choose between *E. coli* and *Comamonas sp.,* a bacterial species previously known to be a preferred food source for wild-type worms (Shtonda et al. 2006). We found that wild-type worms indeed preferentially select *Comamonas* over OP50 *E. coli* (Figure 4C), and that this preference increases over time, reaching a maximum at 24 hours. In contrast, *prp-40* mutant animals do not exhibit this preference and instead choose OP50 over *Comamonas*. These results indicate that mutation of a global regulator of microexon splicing alters food preference behavior.

**Figure 4:**
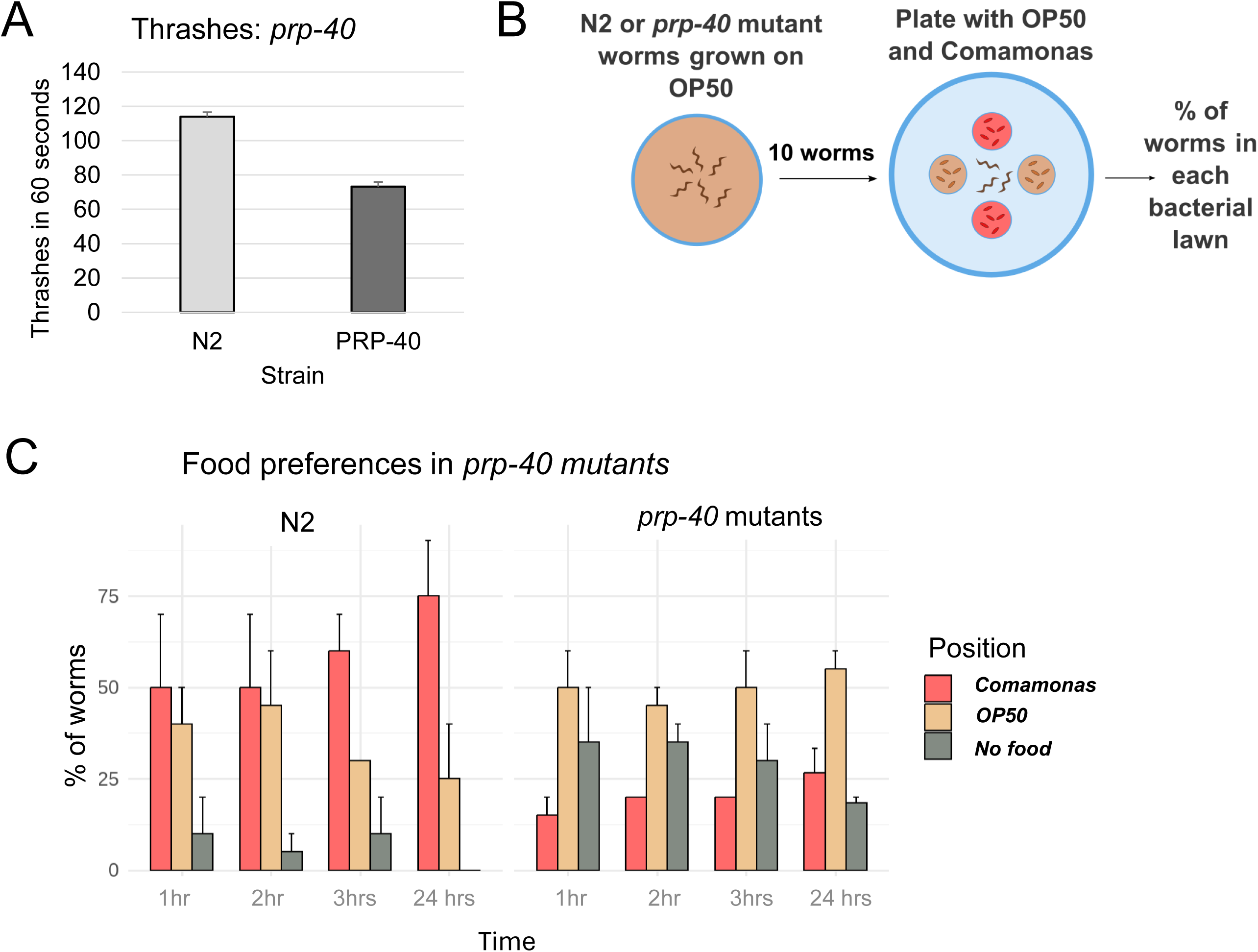
Mutation of the microexon regulator *prp-40* alters food preference behavior. (A) Quantification of the number of body bends (thrashes) per minute. Data represent means ± SD (n =10). (B) Food preference protocol. (C) Percentage of worms that choose each food type for wild type and *prp-40* mutant animals. Pink bars, *Comamonas sp*; orange bars, *E. coli*; gray bars, no food. Data represent means ± SD (n =10).

### A topoisomerase I inhibitor alters the regulation of AS in response to food availability

Next, we performed the food availability experiment in the presence of camptothecin (CPT), an inhibitor of topoisomerase I (TOP1). TOP1 relieves the supercoiling that occurs during transcription by introducing a nick in the DNA, allowing rotation around the TOP1–DNA complex. CPT specifically targets TOP1 and is therefore considered an inhibitor of transcription elongation which in turn affects splicing (Listerman et al. 2006, Dujardin et al. 2014) as it blocks TOP1 activity. Importantly, CPT affects elongation within the gene body rather than at the promoter (Collins et al. 2001).

We found that treating animals with CPT reduced or abolished the effect of food on AS for all three microexon events examined. In the case of *kvs-5*, the effect of food availability on AS is reduced in the presence of CPT (Figure 5A). In the case of *tyra-2*, the tyramine receptor associated with the response to food encounter, the effect on AS is totally abolished in the presence of CPT (Figure 5B). CPT treatment also abolishes the AS response to food availability in *lgc-40*, the serotonin receptor that also mediates feeding responses. (Figure 5C). As a control, *glr-4* is shown, where there are no changes in AS causes neither by food nor by CPT treatment (Figure 5D). These results indicate that inhibition of TOP1 by CPT disrupts microexon AS response to food availability, thus suggesting that this AS response is mediated by TOP1 activity.

**Figure 5:**
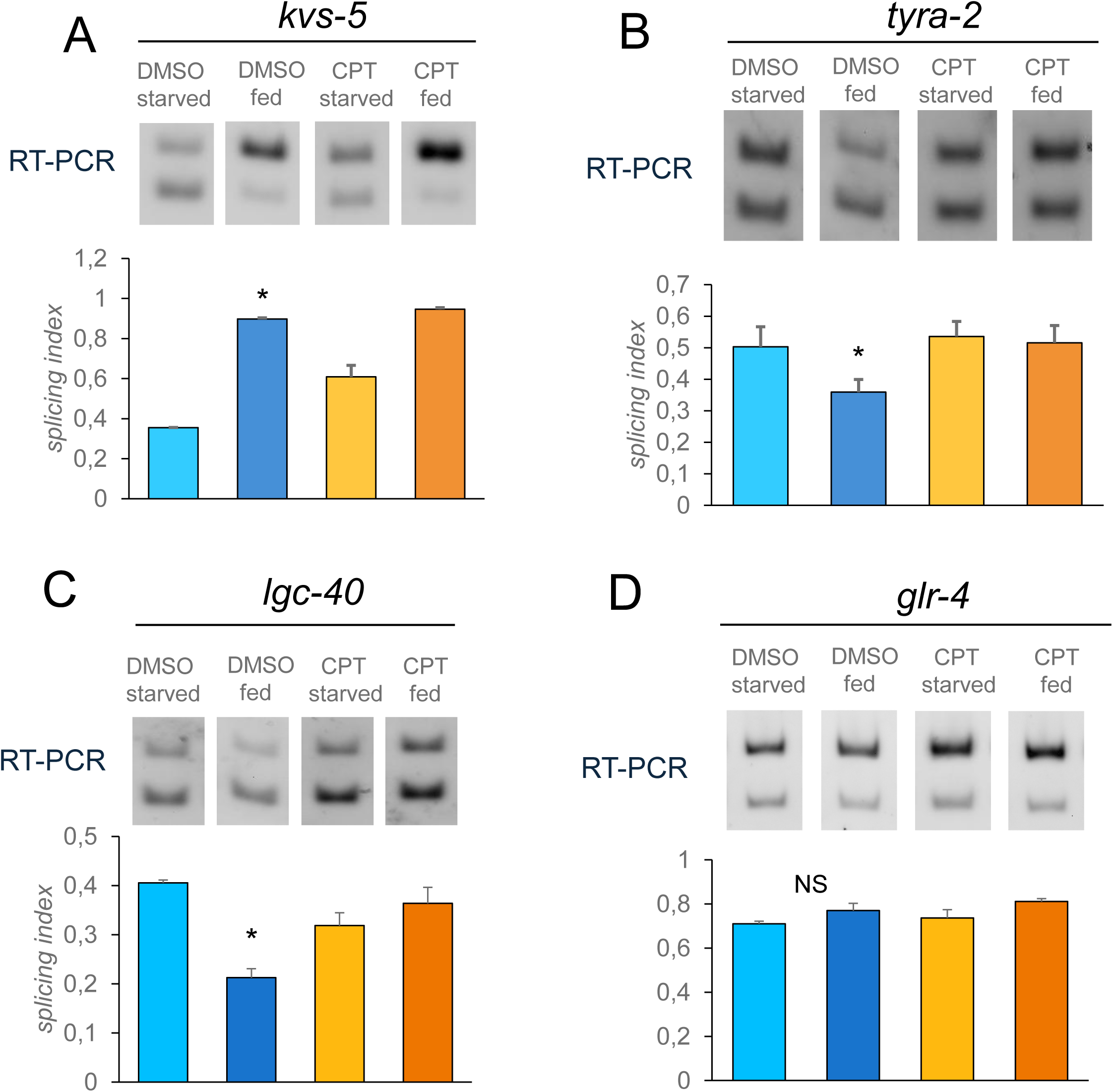
A topoisomerase I inhibitor alters the regulation of AS in response to food availability. Top: Representative RT-PCR gel images for the alternative splicing patterns of *kvs-5* (A), *tyra-2* (B), *lgc-40* (C) and *glr-4* (D). Bottom: quantification of SI. Light blue, DMSO starved worms; dark blue, DMSO starved then fed worms; yellow, starved worms treated with 90 µM CPT; orange, starved then fed worms treated with 90 µM CPT. Data represent means ± SD (n ≥ 3). **\*** p < 0.05 (Student’s t-test).

### CPT modifies the behavioral response to food availability

As we observed that CPT affects microexon AS response upon food availability, we next sought to determine whether CPT could also modify the stereotyped behavior of worms of slowing down locomotion at food encounter.

Before performing the behavioral assay, we first assessed locomotor ability using a thrashing assay in the presence of CPT. We found that CPT did not affect the motor capacity of the animals, which makes our CPT treatment suitable to perform a standard food encounter behavioral assay (Supp. Fig. 2).

In this assay, animals are placed in the center of a Petri dish containing an OP50 *E. coli* lawn—the food—spatially restricted to a ring along the edges (time = 0). Worms were filmed from 5 sec. before to 5 sec. after reaching the bacterial edge. Each worm is recorded individually, and the videos are subsequently analyzed using tracking software. We chose to study four conditions: starved and fed animals in the presence of vehicle, and starved and fed animals in the presence of CPT (Figure 6A). We quantified the animals’ speed before and after encountering the food under control conditions (Figure 6B, left) and before and after encountering food in the presence of CPT (Figure 6B, right). The change in speed upon food encounter is visualized by the statistically significant differences of the curves under the brackets and quantified as a slowing index (Figure 6C).

**Figure 6:**
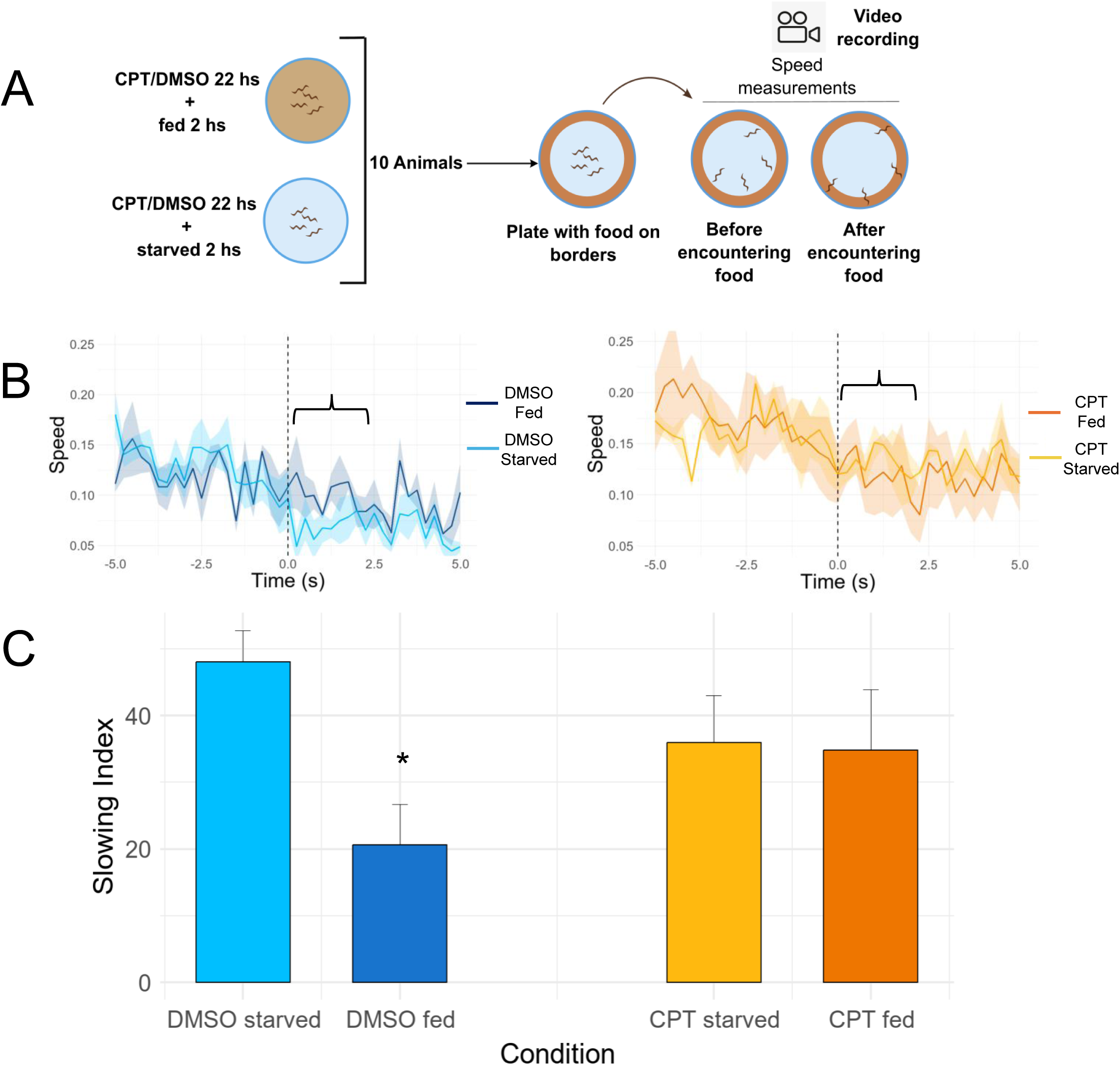
CPT modifies the behavioral response to food availability. (A) Behavioral food encounter protocol. (B, left) Average speed over time for DMSO fed and starved worms and (B, right) average speed over time for CPT fed and starved worms. Time=0 represents the moment of food encounter. The curves are superimposed for easier visualization. Shaded areas represent mean ± SE (n ≥ 4). (C) Quantification of the Slowing Index, defined as the percentage reduction in speed after food encounter relative to the speed before food encounter. Light blue, DMSO starved; dark blue, DMSO fed; yellow, CPT starved; orange, CPT fed. Bars represent mean ± SE (n ≥ 4). *p < 0.05 (Student’s t-test).

As expected based on the behavior described for wild-type worms, starved and fed animals exhibit a reduction in speed upon encountering food. This speed decrease is higher in the case of starved worms. In contrast, this difference in speed decrease between starved and fed worms is completely abolished in the presence of CPT (Figure 6C). These results indicate that CPT treatment disrupts the normal behavioral response to food encounter.

### CPT alters exploratory behavior in response to food availability

To complement the speed assay, we performed a behavioral test focused on spatial exploration rather than speed. In this experiment, animals from four conditions (starved, fed, starved with CPT, and fed with CPT) were individually placed on a food-containing plate marked with a grid and allowed to explore for three hours. After this period, the tracks left by each worm were analyzed, and the number of grid squares was counted (Figure 7A). During this period, the worms can engage in two types of strategies: an active exploratory mode and a sedentary mode (Flavell et al, 2013). A satiated animal spends a greater proportion of time in the exploratory state, traversing a larger number of grid squares on the plate, whereas a starved animal spends more time feeding in the sedentary state, and therefore explores fewer grid squares. (Flavell et al, 2013).

**Figure 7:**
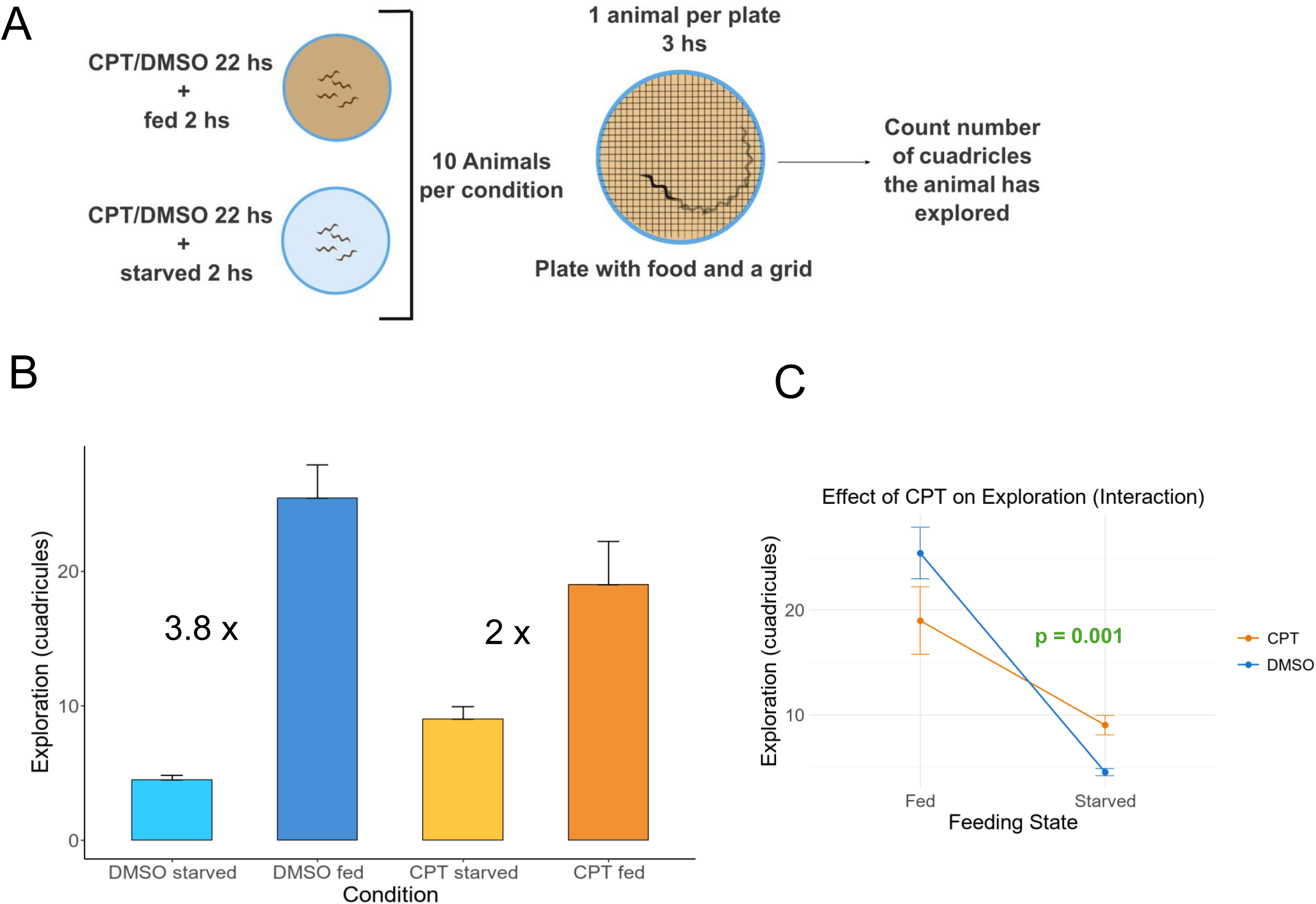
CPT alters exploratory behavior in response to food availability. (A) Exploratory behavior protocol. (B) Quantification of the number of grid squares explored in each condition. Light blue: DMSO starved; dark blue: DMSO fed; yellow: CPT starved; orange: CPT fed. Bars represent mean ± SE (n ≥ 9). Fold changes between starved and fed conditions are shown. (C) Interaction plot depicting the differences in grid squares explored in starved and fed conditions. Blue, treatment with DMSO; orange, treatment with CPT 90 μM. Dots represent mean ± SE (n ≥ 9). Interaction between nutritional states and treatment with CPT was evaluated with 2-factor ANOVA (p < 0.001).

Quantitative analysis of these data is presented in Figures 7B. Under control conditions, fed animals explored nearly four times as many squares as starved animals. In the presence of CPT, this difference was reduced to approximately twofold. Furthermore, we performed a two-factor ANOVA and there is a significative interaction between feeding state and CPT treatment (Figure 7C). These results indicate that CPT modulates exploratory behavior in response to food availability.

As a first attempt to investigate a possible mechanism by which microexon AS promotes changes in speed reduction and exploratory behaviors in response to feeding, we decided to look at PRP40 mRNA expression levels and found that they do not change upon starved and fed animals, nor upon CPT treatment (Supp. Fig. 3). This suggests that, although both PRP-40 and CPT modulate microexon AS and feeding behavior, they seem to act through independent mechanisms.

## Discussion

In this study we show that microexon AS is regulated between *C. elegans* larvae and adults (Figure 1). Furthermore, microexon inclusion is differentially regulated among distinct neuronal types (Figure 2), which shows that microexon splicing is spatially regulated within the nervous system. Food availability, an environmental stimulus, regulates microexon inclusion of a subset of genes including *tyra-2* and *lgc-40,* which are important receptors in food-related behaviors (Figure 3). In addition, CPT, a topoisomerase I inhibitor, disrupts this regulation of AS in response to food availability (Figure 5).

We provide evidence to support that CPT treatment alters both *C. elegans* microexon AS and behavioral responses to food availability. Regarding animal behavior, we observe both changes in locomotion speed reduction and in exploratory behavior (Figures 6 and 7). Importantly, mutation of the microexon regulator *prp-40* alters food preference behavior (Figure 4). However, although we cannot conclude that microexon AS causes the observed changes in speed reduction and exploration behaviors since, as explained, this experiment cannot be performed with *prp-40 mutant* animals as their locomotion is heavily affected, altogether our results suggest that microexons are involved in animal behavior.

Our observations represent an initial step towards exploring the physiological consequences of AS regulation in response to environmental stimuli. In a previous study, we investigated how plants modulate transcriptional elongation in response to light and darkness, which in turn leads to changes in alternative splicing. This regulation occurs at the level of the whole organism (Godoy Herz et al. 2019). In the present study, we aimed to examine the consequences of AS regulation also in a whole organism, this time in an animal, in which the added dimensions of locomotion and behavior introduce further complexity to the question.

To investigate the role of microexons in detail, future studies will need to focus on the function of individual microexons, as exemplified by recent work such as Choudhary et al. 2025 in *C. elegans* and studies conducted in the zebrafish model (Calhoun et al. 2025).

Briefly, Calhourn and colleagues used CRISPR/Cas9 to remove individual microexons in zebrafish and assessed larval brain activity, morphology, and behavior. Surprisingly, most mutants had minimal or no phenotypes. Such approaches are inherently time-consuming, as they require the generation of transgenic animals. Moreover, phenotypic effects are often difficult to detect— as observed in the zebrafish study—likely due to genetic and functional redundancy, which can mask observable phenotypes.

Regarding the effects of CPT, this compound and its derivatives are widely used in antitumor therapies (Takimoto et al. 1998). Its sole molecular target is TOP1. TOP1 relaxes DNA supercoiling in the absence of an energy cofactor by introducing a nick in the DNA and allowing the cleaved strand to rotate around the TOP1-bound complementary strand. TOP1 is the only target of CPT (Pommier et al. 2006). TOP1 is required for transcriptional elongation because it removes the supercoiling generated by elongation complexes. In addition, it has been reported that TOP1 can phosphorylate certain SR proteins—thereby influencing alternative splicing—and can also associate with TFIID to repress transcription initiation (Collins 2001, Pommier 2006, Khobta 2006). About the mechanism of action of CPT: when camptothecin freezes the TOP1–DNA complex, the enzyme is unable to complete the reaction cycle. The topological state of the domain encompassing the frozen TOP1 remains fixed until the enzyme is liberated from CPT (Khobta et al. 2006). Therefore, CPT treatment inhibits transcriptional elongation.

Our results demonstrate that inhibition of TOP1 by CPT disrupts microexon AS response to food availability. This indicates that the mechanism of microexon AS regulation is influenced by TOP1 activity. Interestingly, it has been reported that topoisomerases are mutated in some individuals with ASD. Furthermore, inhibition of TOP1 in mouse and human neurons demonstrates that TOP1 is necessary for transcription of long genes related to ASD. This effect on gene expression is due to impaired transcription elongation (King et al. 2013). Since microexons are also strongly associated to ASD (Irimia et al. 2014), perhaps, in addition, TOP1 could play a role in regulating microexon AS.

If microexon AS is impaired, as shown in *prp-40* mutant animals, we observe changes in food preference behavior (Figure 4). Thus, microexon AS produces changes in this particular behavior. On the other hand, CPT treatment shows that TOP1 is involved in the regulation of microexon AS and that this is, in turn, correlates with alterations in speed reduction and exploratory behaviors in response to food availability. Our results are consistent with previously reported findings showing changes in AS between starved and fed L1 larvae. In the absence of food, L1 arrest their development. This coincides with RNA Polymerase II accumulation at promoters of certain genes (Baugh et al. 2009). Although this protocol is more severe because it alters the animal’s development, it shows that nutritional changes can alter transcription elongation, similar to our findings that inhibition of TOP1 alters the regulation of AS in response to food availability.

Due to its complexity, the search for the mechanism that shows how microexons are involved in behavioral plasticity exceeds the present report and future work from different groups will be needed to further investigate the consequences of AS regulation in whole organisms. Together, these findings demonstrate that the disruption of microexon AS can have consequences on both development and animal behavior, highlighting a clear link between environmental sensing, splicing regulation, DNA topology and behavioral output.

## Materials and methods

### Animals and growth conditions

All animals were maintained at 20°C on nematode growth media (NGM) agar plates with the OP50 *E. coli* strain as a food source. Experiments were performed on young adults unless otherwise specified. Bristol N2 was used as a wild-type strain. *prp-40* mutants were provided by Adam Norris (strain: *prp-40(csb3)/hT2)*.

### RT-PCR for alternative splicing assessment

Total RNA extraction of worms was carried out using the TRI-Reagent (MRC) following the manufacturer’s instructions. 500 ng of RNA were further used to synthesize cDNA with MMLV-RT and oligo-dT as primer following the manufacturer’s instructions. cDNA was amplified via PCR using primers mapping in the exons that flank the microexons, to amplify both the microexon including and excluding isoforms. PCR products were analyzed with 6% polyacrylamide gel electrophoresis and the quantification was done on Image J.

### Neuron-specific microexon analysis

Analysis of neuron-specific microexon alternative splicing was performed using CeNGEN single-cell transcriptomes (Taylor et al, 2021; Weinreb et al, 2025). We used the available mPSI (mean Percent Selected Index) data, which quantifies splice junction usage, for each neuronal type. Mean and standard deviation are shown. In the case of *tyra-2* and *kvs-5*, all neurons with available data are presented. In the case of *uaf-1* and *unc-2* only the data of the neurons shared between both events are shown. To assess whether the variances of mPSI between each microexon and the control exon *uaf-1* differ significantly, we performed a homoscedasticity test (Levene’s test). For each comparison, we used data from all shared neuron types.

Neuron types: PVC (Posterior Ventral Process C neurons), PVQ (Posterior Ventral Process Q neurons), SIA (Sublateral Interneurons A), DVB (Dorsorectal Ganglion Ventral Process B neuron), SMD (Sublateral Motor neurons D), VD (Ventral D-type Motor Neurons), CEP (CEphalic Sensory neurons), OLL (Outer Labial Lateral neurons), ADF (Amphid Dual Ciliated Ending F neurons), AFD (Amphid Finger-like Endings D neurons), ASER (Amphid Single Cilium E Right neuron), ASG (Amphid Single Cilium G neurons), ASI (Amphid Single Cilium I neurons), ASK (Amphid Single Cilium K neurons), AWA (Amphid Wing A neurons), AWC (Amphid Wing C neurons), PHA (PHasmid A neurons), AIM (Anterior Interneurons M), AVH (Anterior Ventral Process H neurons), DVC (Dorsorectal Ganglion Ventral Process C neuron), RIC (Ring Interneurons C), RIM (Ring Interneurons M), RMD (Ring Motor neurons D), RME (Ring Motor neurons E), AVM (Anterior Ventral Microtubule neuron), IL1 (Inner Labial 1 neurons), ADL (Amphid Dual Ciliated Ending L neurons).

### Food availability protocol

Young adult N2 worms were transferred to NGM agar plates without a bacterial lawn and maintained for 2 h to induce starvation. Worms were then either transferred to NGM plates containing a fresh *E. coli* lawn (fed condition) or to new NGM plates without a lawn (starved condition) for an additional 2 h. Following these treatments, total RNA was extracted. Copper wire rings were placed on all plates to prevent starved worms from climbing the plate walls (Davies et al. 2003).

### Drug treatments

L4 worms were transferred to 90 µM CPT or 0,36% v/v DMSO plates for 22 h. Then, worms were transferred to NGM plates with CPT or DMSO without a bacterial lawn for 2 h to induce starvation. This completes the 24-hour incubation with CPT. Then, worms were either transferred to NGM plates containing a fresh *E. coli* lawn (fed condition) or to new NGM plates without a lawn (starved condition) for additional 2 h. For alternative splicing assessment, total RNA was extracted. See below for behavioral assays.

## Behavioral assays

### Thrashing assay

L3 stage animals of respective genotypes (N2 and *prp-40(csb5)*) were transferred on 30 µL M9 buffer droplets on unseeded plates and the number of thrashes were counted for 60 s. The same assay was done to worms treated with CPT or DMSO.

### Food preference assay

NGM plates were seeded with 2 droplets of OP50 *E. coli* and *Comamonas sp.* bacteria fresh cultures, and were left to grow overnight. To start the assay, N2 and *prp-40* mutant L3 worms were placed on the center of the plates, equidistant from each of the 4 droplets (1 cm away). Worms’ positions were observed each hour for 3 hours, and the animals were left overnight at 20°C to be measured 24 hours later.

### Slowing Response Assay

After treatment with CPT/presence or absence of food, a droplet of M9 buffer containing adult worms was placed at the center of NGM plates with food only on their borders, and was allowed to dry before recording. Locomotion was captured at 30 frames per second using an Allied Vision Technology Guppy Pro GPF 125C IRF Camera. Locomotion velocity was quantified using the Multi-Worm Tracker (Rex Kerr, https://sourceforge.net/projects/mwt/) and analyzed using custom MATLAB scripts. To assess the food-induced slowing response, we compared the speed upon encountering the bacterial lawn (up to 5 seconds after) and the speed just before encountering it (up to 5 seconds before).

### Exploration Assay

To measure exploration behavior, after treatment, individual adult worms were placed in NGM agar plates uniformly covered in bacterial lawn. After 3 hs, the worms were removed and a grid containing 5 mm squares was superimposed on the now empty plates. The number of squares entered by the worm’s tracks was counted manually (maximum 106 squares).

## Acknowledgments

We thank María Eugenia Martín, Valeria Buggiano and Luisa Cochella for their invaluable help. This work was supported by grants from the Agencia Nacional de Promoción de la Investigación, el Desarrollo Tecnológico y la Innovación of Argentina (Agencia I+D+i) PICT-2019-03787, the Consejo Nacional de Investigaciones Científicas y Técnicas of Argentina (CONICET) PIBAA-2022 and the Lounsbery Foundation. M.A.G.H., D.R. and A.R.K. are career investigators from CONICET. G.M. received a fellowship from Agencia I+D+i.

