## Supplemental Figure 3 for "Microexon alternative splicing and feeding behavior in *C. elegans*"

*Suppl. Fig. 3*

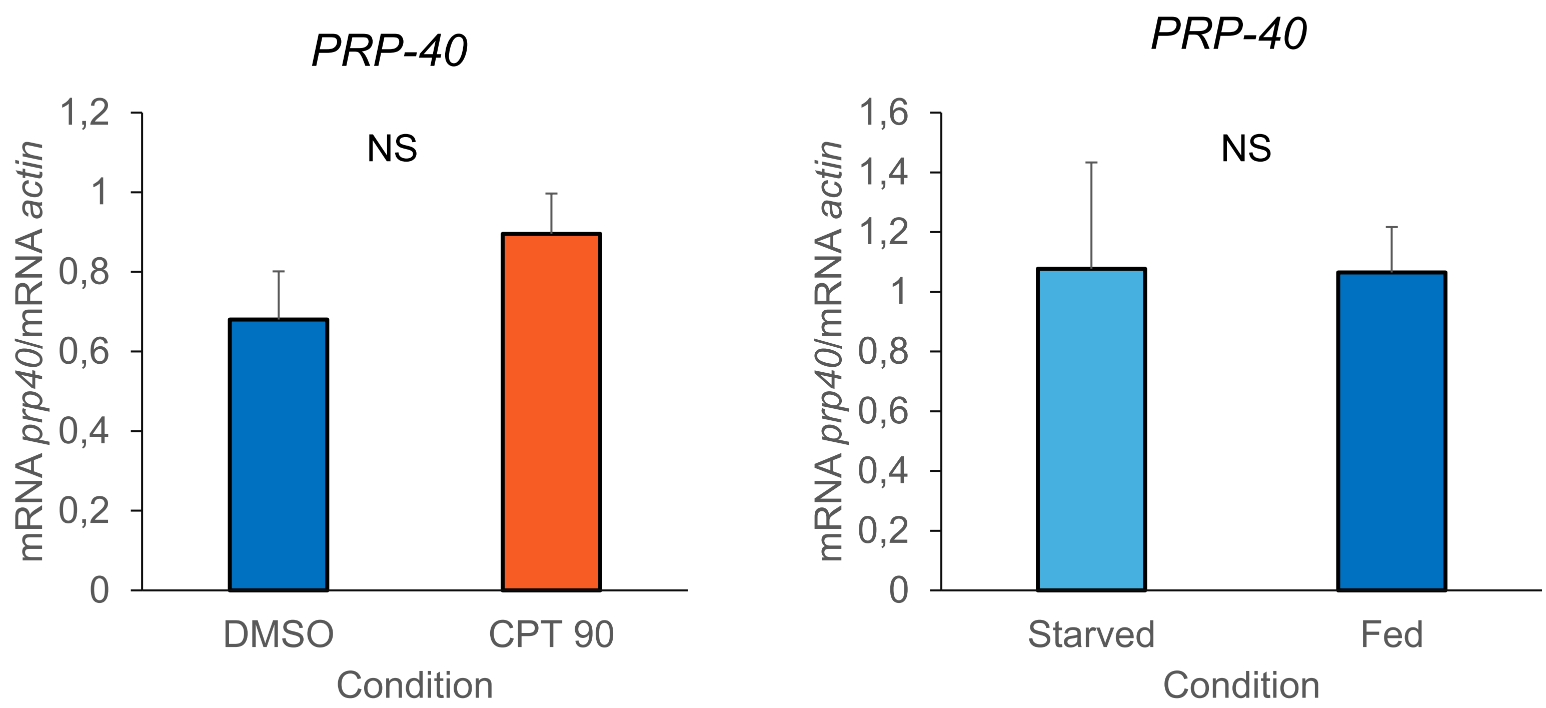


**Suppl. Fig. 3: PRP40 expression levels.** PRP40 relative to actin mRNA measured by qPCR in DMSO and CPT treated animals (**A**) and starved and fed animals (**B**). Data represent means ± standard deviation ($n \geq 3$).
