## Supplemental Figure 1 for "Microexon alternative splicing and feeding behavior in *C. elegans*"

*Suppl. Fig. 1*

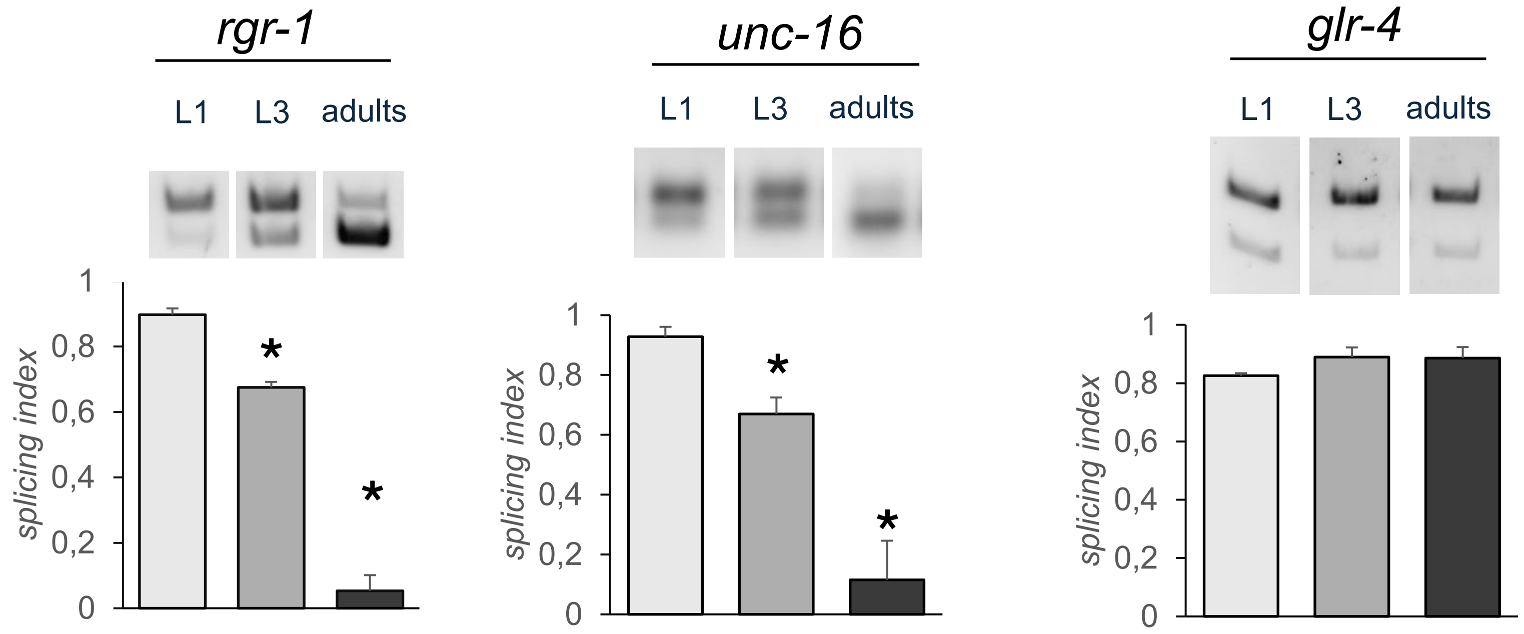


**Suppl. Fig. 1: Microexon AS changes throughout *C. elegans* development in L1, L3 and adult samples**. Related to Figure 1. Top: representative RT-PCR gel images for the alternative splicing patterns of *unc-13, rgr-1, tyra-2, nmy-1, unc-16* and *glr4*. Bottom: quantification of splicing index. White bars, L1; gray bars, L3; dark gray bars, adults. Data represent means ± standard deviation (n ≥ 3).
