## Supplemental Figure 2 for "Microexon alternative splicing and feeding behavior in *C. elegans*"

*Suppl. Fig. 2*

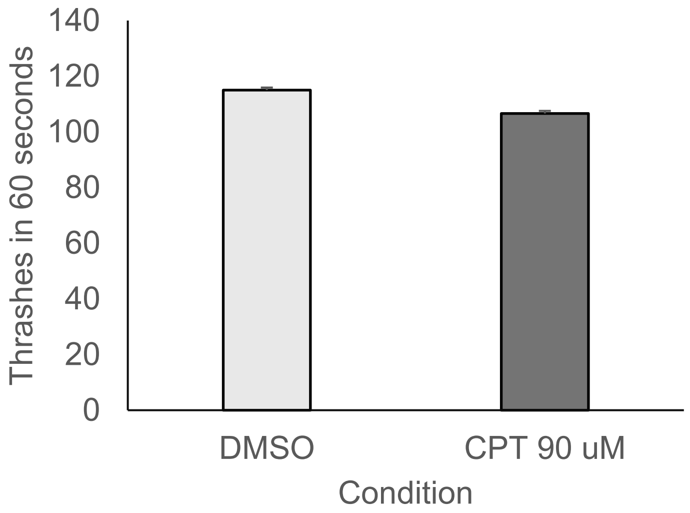


**Suppl. Fig. 2: CPT treatment does not affect locomotion**. Animals were treated with DMSO or CPT 90 μM. Quantification of the number of body bends (thrashes) per 60 seconds. Data represent means ± standard deviation (n ≥ 3).
